# Long-read sequencing to detect full-length protein-protein interactions

**DOI:** 10.1101/2024.04.01.586447

**Authors:** Stephanie Schaefer-Ramadan, Yue Guan, Ayeda A. Ahmed, Jovana Aleksic, Khadija A. Elmagarmid, Leena F. Syed, Yasmin A. Mohamoud, Joel A. Malek

## Abstract

Given the increased predictions on interactome size and demand for protein function information, methods for detecting protein-protein interactions remain a significant development area. The all-vs.-all sequencing (AVA-Seq) method utilizes a convergent fusion plasmid design to make two-hybrid technology amenable to next-generation sequencing. Here, we further innovate to take advantage of synthetic DNA technologies and Oxford Nanopore Technologies long-read sequencing improvements to allow us to determine full-length protein-protein interactions. Here, using this approach we recovered 159 protein-protein interactions from a set of 57 human proteins using multiple forms of validation. Further, when referencing a human gold standard set of interactions, eight full-length protein-protein interactions were recovered from an expected 28 interaction pairs (28.6%), a typical recovery rate for two-hybrid technologies. The AVA-Seq, in combination with the ease of synthetic DNA production and the MinION platform, offers a low-cost, high-throughput alternative for determining protein-protein interactions, which can be utilized in research labs at all stages.

**Key Points:**

1. First application of long-read sequencing for full-length protein-protein interaction studies.
2. The recovery rate of the AVA-Seq method using full-length proteins is on par with other leading methods.
3. Advances in synthetic biology and sequencing technologies make full-length protein interactomes affordable and accessible.

**GRAPHICAL ABSTRACT:** 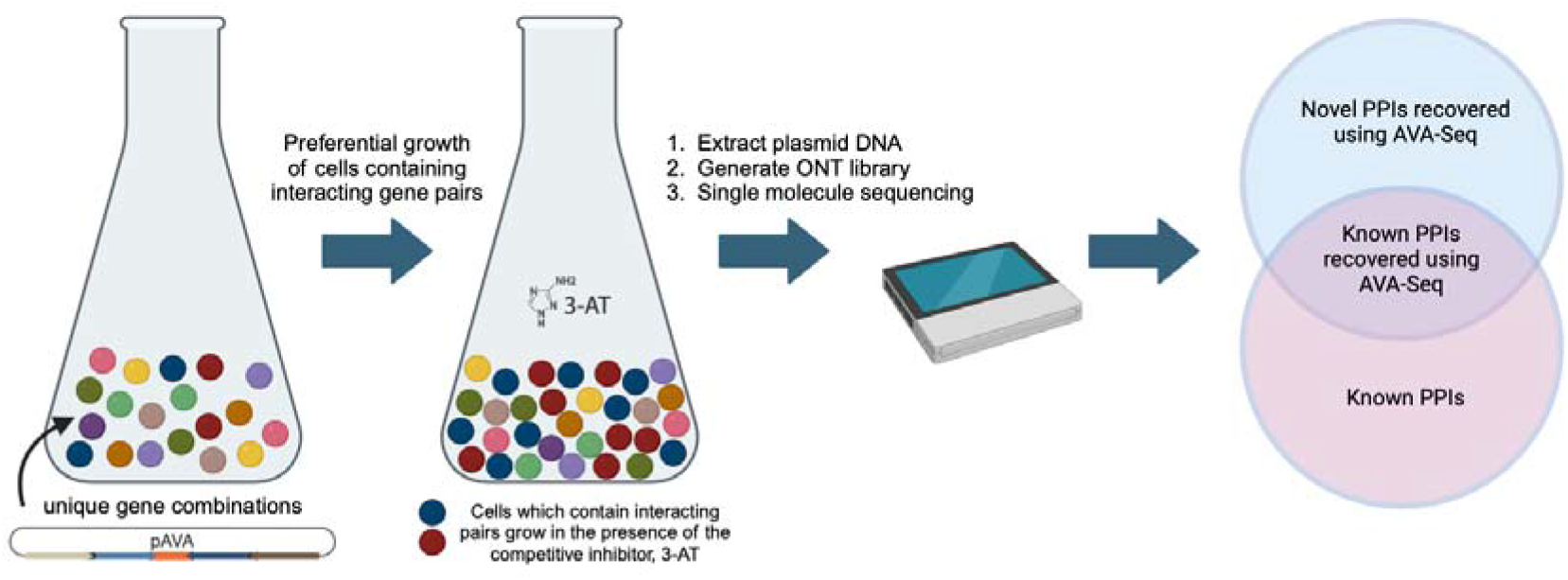

## INTRODUCTION

Methods for protein interaction mapping that utilize the cost and speed benefits from next-generation sequencing (NGS) are limited (1–10). In the case of two-hybrid systems, this is primarily because the information about which ‘bait’ and ‘prey’ interact are often on separate plasmids that are not physically connected. In recent years, our lab has designed a convergent fusion plasmid (pAVA), allowing for high-throughput sequencing of interacting protein fragments in a bacterial system (8). This novel concept enables users to utilize the bacterial two-hybrid system while simultaneously exploiting liquid culture and NGS platforms for significant speed improvements. Until recently, we exclusively used gene fragments in the range of 360-450 bp (120 to 150 amino acids) for a combined size of ∼800-900 bp, the amplicon size limit for most short-read technologies. While the information derived from protein fragments of this size can provide information on the broader protein interaction, it would be beneficial to develop the all-vs.-all sequencing (AVA-Seq) system to allow full-length protein testing.

Bacterial two-hybrids are valuable tools in the protein interaction field and are robust in determining known and novel protein-protein interactions (PPIs) across many species. Recently, full-length SARS-Covid ORFs were utilized in a bacterial two-hybrid system to determine interactions with the human ACE2 protein (11). This application demonstrates that the bacterial system can handle full-length protein interaction studies and could be adapted to third-generation sequencing technologies such as Oxford Nanopore.

In this work, we utilized a subset of proteins from the gold-standard for protein interactions, the human positive reference set (HsPRS-V2 (12); referred to as PRS), which could be synthesized on a single DNA fragment (less than 1.8 kb each). The goal was to adapt the AVA-Seq method to accommodate full-length proteins, allowing a more direct comparison to traditional protein interaction methods.

The long-read sequencing technology by Oxford Nanopore Technologies (ONT) allows for novel genome assembly (13), improving existing genomes (14, 15), epigenetic studies (16), and transcriptome splicing analysis, among other applications that benefit from longer sequences. The ONT platform has also been utilized for interactions between RNA binding proteins with RNA (17), protein-DNA interactions (18–20), host-virus interactions (21), host-pathogen (22) (plant gene-for-gene network), and enhancer-promoter interactions (23). To our knowledge, single-molecule, long-read sequencing has not yet been applied to studying protein-protein interactions. This study demonstrates the combination of AVA-Seq convergent fusion design with the long-read capabilities of ONT to provide a rapid, low-cost protein interactome.

## MATERIAL AND METHODS

### Pooling of proteins

Synthetic genes were codon optimized for E. coli and contained specific 5’- and 3’-adapter sequences to facilitate directional PCR amplification. 1 µg of lyophilized DNA were resuspended in 50 µL of ultrapure water. Genes were pooled at equal molar concentrations. Two different gene pools were made, reflecting different gene lengths. The short gene pool (genes <1,299 bp) contained 35 genes plus two additional frameshifted fragments. The long gene pool (genes >1,300 bp) contained 22 genes plus two additional long frameshifted fragments. The frame-shifted fragments code for an early stop codon and help assess potential auto-activators. The protein pools were used the same day for PCR amplification and stored at −20 °C.

### Generating the full-length protein-protein fusions

DNA was synthesized (TWIST Bioscience) for 57 human genes using codon optimization for expression in E. coli, and specific 5’- and 3’-adapter sequences were added. These adapters allow for directional amplification using different overhanging primers, allowing two genes to be merged into the convergent fusion DNA strand. After assembly, the inserts have an equal chance of being fused to the DBD or the AD contained in the pAVA plasmid. This also allows for simultaneous screening in multiple orientations. Two separate amplifications were performed using the pooled DNA to ensure high coverage of all proteins.

### Ligation into pAVA plasmid and preparation for 3-AT selection

A ligation ratio of 1:1 vector to insert was used to reduce multiple insertions. The column-cleaned inserts are ligated using T4 DNA ligase (NEB M0202) to the linearized pAVA plasmid [DPN1 (NEB R0176) and rSAP (NEB M0371) treated before use] and incubated at 16 °C overnight. The ligation product is column-cleaned (Sigma-Aldrich NA1020) and transformed using NEB Turbo Electrocompetent cells (NEB C2986; discontinued). The DNA were then extracted (Sigma-Aldrich PLN350) and transformed using the validation reporter cell line (24).

### 3-AT selection and controls

3-AT interaction selections were performed using the transformed validation reporter cell line as previously reported (25). High-throughput AVA-Seq validation: experiments were conducted in triplicate. Each experiment contained three technical replicates of 0 mM, 2 mM, and 5 mM 3-AT liquid growth conditions at 37 °C for 9 hours, as reported previously (8, 25, 26).

Low-throughput validation: For the individual full-length PRS protein pairs, cultures were washed four times in minimal media and diluted to a final concentration of OD_600_ = 0.015. Constructs were screened in a 96-deepwell plate with a volume of 1 mL in quadruplicate for all three conditions (0 mM, 2 mM, and 5 mM 3-AT) at 37 °C for 9 or 16 hours. The OD_600_ of the endpoint was measured using a standard 96-well plate with 100 µL of minimal media and 40 µL of culture using the minimal media as a blank. Measurements were normalized to the final 0 mM readings.

### MinION library preparation and sequencing

The DNA were extracted from each 3-AT replicate sample (Sigma-Aldrich PLN350) and quantified using QuBit HS (Invitrogen Q32854). Three 0 mM, three 2 mM and three 5 mM samples make up one experiment. Twenty-seven libraries (9 for each replicate experiment) were prepared using the Ligation Sequencing Kit protocol for Ligation sequencing amplicons - native barcoding (SQK-LSK109 with EXP-NBD196, Oxford Nanopore Technologies). Briefly, 130 ng for 1 kb amplicon DNA was prepared for each sample. Then amplicon DNA were end-repaired using NEBNext Ultra II End-Repair/dA-tailing Module (NEB E7546). Subsequently, each sample was cleaned with AMPure XP beads (Beckman Coulter A63881). Ligation of the Native barcode was then performed by adding Blunt/TA Ligase Master Mix (NEB M0367) and a unique native barcode (Oxford Nanopore Technologies EXP-NBD196) for each end-prepped DNA. The samples were then cleaned with AMPure XP beads and quantified with QuBit HS. The library pool was prepared by combining equimolar amounts of barcoded samples into an Eppendorf tube for a final molarity of 0.4 pmoles. Finally, the adapter ligation and clean-up were performed following the manufacturer’s instructions. To ensure all DNA fragments are retained, the pool was cleaned using Short Fragment Buffer (Oxford Nanopore Technologies SQK-LSK109). The resulting pool ∼80 fmol was then loaded onto the MinION flow cell (Oxford Nanopore Technologies FLO-MIN106D) and sequenced for 72 hrs.

### Data analysis

FASTQ sequence files from the Oxford Nanopore sequencer contain long reads spanning the two encoded convergent proteins being tested. The FASTQ files were searched for the multiple STOP insert sequence (8) using custom Perl scripts allowing for fuzzy sequence matching. Sequences to the right and left of the STOP insert were retained and searched for sequences indicating a fusion to either the lambda CI (DNA binding domain; DBD) or RNAP (activation domain; AD) of the system. Both sequences were then searched against the sequence database for this project using BLASTN (27). Paired sequences that exhibited high quality BLAST matches with our sequence database were retained for further analysis. Subsequently, protein pairs were compiled, and each instance of identical protein pairs observed across separate read pairs led to an increment in count. The counts for each protein pair across all nine replicates (comprising triplicates of 0 mM, 2 mM, and 5 mM 3-AT) were aggregated into a table for subsequent statistical analysis in R.

Protein pairs were removed if they had a cumulative count of less than 10 reads in the combined three baseline replicates (0 mM 3-AT). Pairs failing to meet this threshold were excluded due to challenges in statistical processing arising from low count numbers.

Using the R package edgeR (28), protein pairs exhibiting a statistically significant increase under selective conditions (2 mM or 5 mM 3-AT) compared to the background (0 mM 3-AT) were identified. Internally, edgeR performed normalization of count values to accommodate varying sequencing depths reflected in differing library sizes. A negative binomial model was applied to ascertain differential growth, utilizing Fisher’s exact test for significance testing, which computed adjusted p-values or false discovery rates (FDRs) for each protein pair.

Further scrutiny led to the identification of protein pairs meeting specific criteria: a log_2_ fold change (log_2_FC) > 0.8 and FDR < 0.1 in the presence of 3-AT compared to 0 mM 3-AT, denoting potential interactions. Additionally, more stringent filters were applied to eliminate interactions lacking robust support. For the all-versus-all analysis, reporting a PPI necessitated repeatability: at least two experiments per protein pair orientation with log_2_FC > 0.8 and FDR < 0.1. In instances where one orientation exhibited interaction across multiple experiments, the presence of an interaction in the other orientation (with log_2_FC > 0.8 and FDR < 0.1) was acknowledged, even if detected in only one experiment.

## RESULTS

### Individually tested full-length pairs in pAVA (low-throughput)

We first wanted to establish if the convergent fusion design of the pAVA plasmid is usable for screening for full-length protein-protein interactions, as it was initially optimized for shorter fragments. To do this, a selected number of positive reference set (PRS) pairs and random reference set (RRS) pairs (12) were convergently fused and inserted into pAVA in both orientations (fused to DNA binding domain (DBD) or RNAp activation domain (AD)) individually and confirmed by Sanger sequencing.

We performed two initial growth assays. The first was on solid agar (colony formation as a readout), and the second was in liquid media (OD_600_ as a readout) in the absence or presence of 3-AT, a competitive inhibitor of the HIS3 reporter gene. The RRS pairs did not show colony formation on the solid agar in the presence of 3-AT (not shown). Further, all the PRS and RRS pairs we tested had some level of growth in the liquid culture assay.

As we incorporated full-length protein pairs into the pAVA plasmid, we wanted to determine whether the orientation of the genes played a significant role in the growth and resulting observed interaction. There is an equal chance for a given protein-protein pair to show growth when adjacent to either DBD or AD. The PRS pairs tested manually (in the low-throughput liquid growth assay) show growth in at least one orientation as shown in Fig. 1. We observed a preference for some pairs in one orientation versus the other while other pairs grew equally well fused to DBD or AD. Additionally, all positive reference pairs tested showed higher average growth in the mild 2 mM 3-AT growth conditions compared to the strong competitive inhibition of 5 mM 3-AT except for PSMD4|RAD23A (one orientation).

**Figure 1:**
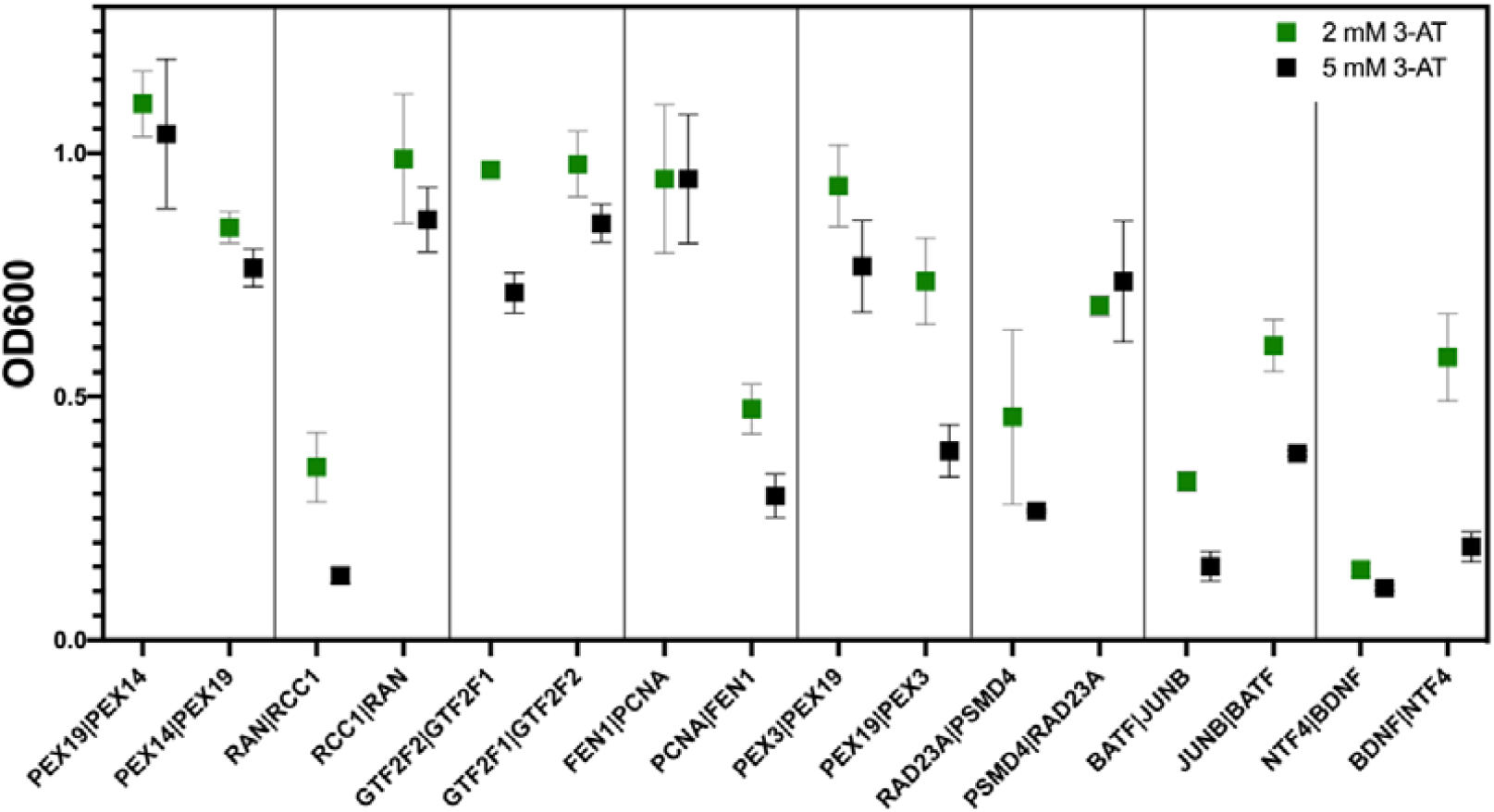
Growth comparison of full-length PRS pairs in both orientations. Data normalized to 0 mM 3-AT 9-hours growth time. The 2 mM growth is shown in green and 5 mM growth is shown in black with error bars (SD). The gene listed first is the DBD fusion.

### Low-throughput assessment of growth time

Due to the large span of protein pair lengths, we wanted to ensure the growth time was sufficient to achieve separation between the 2 mM and 5 mM conditions in all possible protein lengths. To test this, we compared 9- and 16-hour growth in 0 mM, 2 mM, and 5 mM 3-AT conditions for pairs representing different protein-pair lengths. We observed no apparent bias regarding protein length influencing how well the pair grew in liquid media. Our results indicate that a 9-hour growth time is likely sufficient and might be preferred as it gives a more separation in the 2 and 5 mM samples (Supplemental Fig. 1). Furthermore, the 9-hour growth is consistent with our previous AVA-Seq studies (8, 25, 26, 29).

### High-throughput AVA-Seq project design

A protein interaction screening of this dynamic range in length was never attempted previously; therefore, it was difficult to benchmark. Figure 2 outlines our method design. Full-length genes were separated into two sizes to assist with the size selection of the NGS library. This was done for two main reasons. First, this study used a diverse range of gene lengths, and second, the first-generation ONT chemistries preferentially sequence shorter products. As the ONT chemistries improve, testing a true all-vs.-all on this dynamic range of gene products would be worth testing.

**Figure 2:**
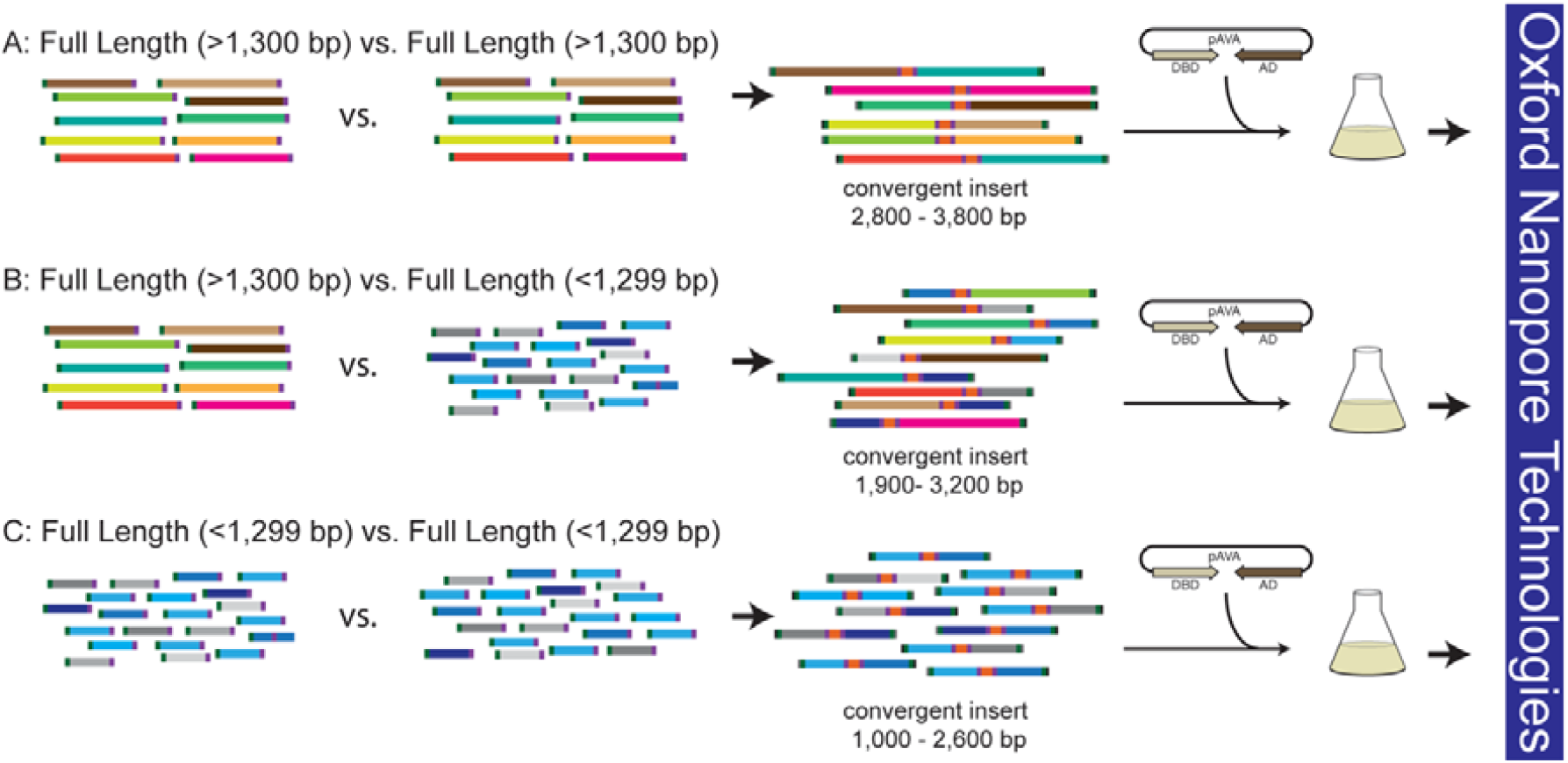
AVA-Seq method experimental design using full-length proteins. A) full-length proteins that are >1300 bp paired and screened against themselves. B) full-length proteins from human proteins >1300 bp paired with full-length proteins <1299 bp. C) full-length proteins <1299 bp paired and screened against themselves. All convergent insert sizes reflect the final DNA length (including adapters) amplified for NGS. Individual protein sequences were synthesized by TWIST Biosciences and included custom adapters added to the 5’ and 3’ ends of the design process. All pathways utilize two sets of primers. Each set has a complementary stop insert, allowing the two fragments to be ‘stitched’ together (represented as orange on the convergent insert).

### Coverage of interaction space (high-throughput)

AVA-Seq can simultaneously screen protein-protein pairs in two orientations, which is advantageous as some fusion pairs can show preferential interaction (12). We obtained over 100x coverage for the gene combinations being tested. These numbers are based on the number of transformants observed as colonies on counting plates used for 3-AT screening events. When considering each orientation as a unique test pair, 3,115 protein-protein pairs were covered out of the possible test space of 3,192 (97.6%). Our requirement for ‘tested’ is the protein pair be in two or more experiments in at least one orientation (3,090 protein-protein pairs). Furthermore, if the first orientation is considered tested and the second orientation is only found in one experiment, it is considered tested in both orientations (25 protein-protein pairs). Figure 3 shows the tested combinations as a heat map.

**Figure 3.**
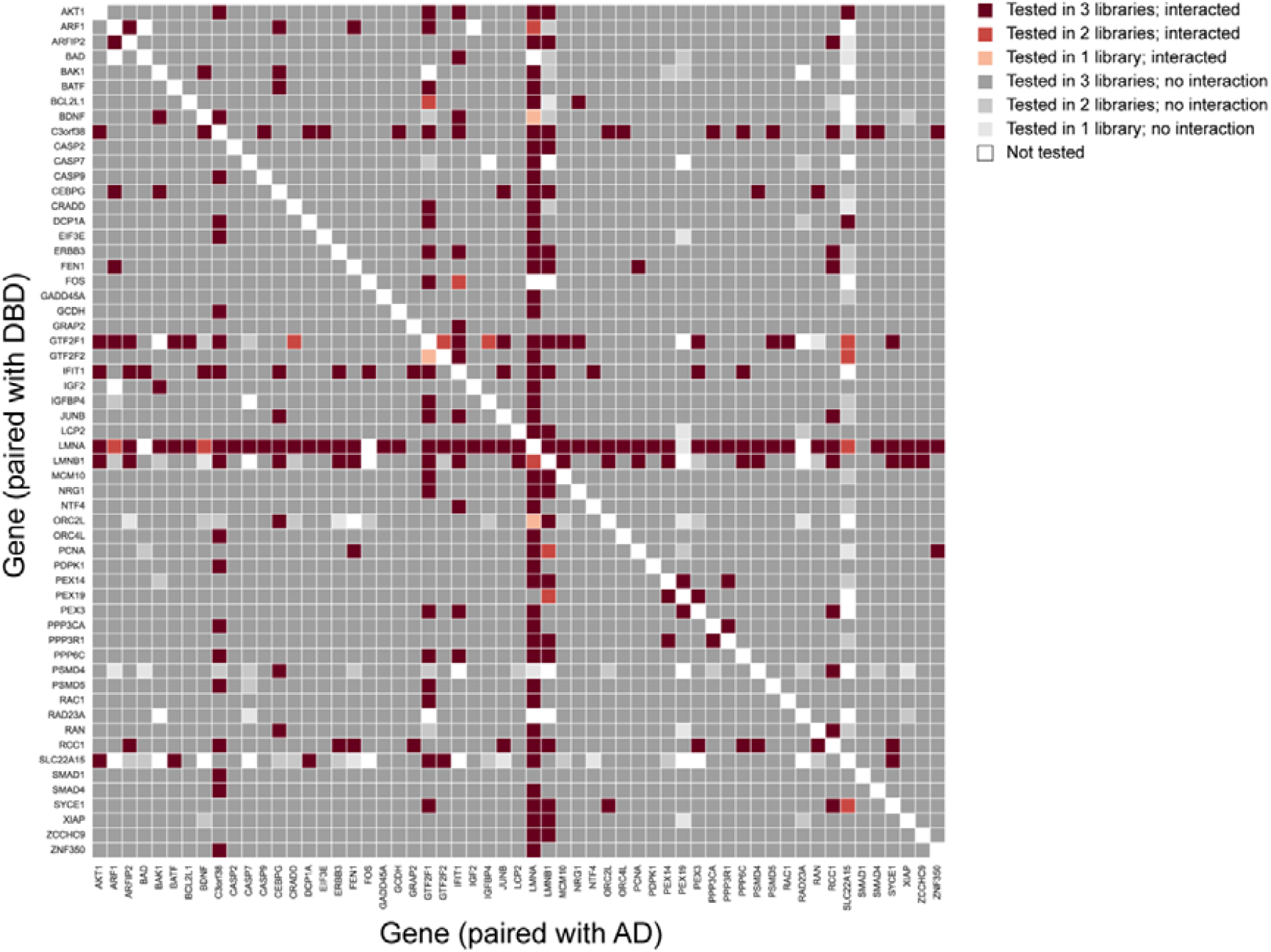
Heatmap of coverage and recovered interactions. The gene pairs tested but did not interact in 3 experiments are colored dark grey and 2 experiments medium grey. The gene pairs tested and determined to interact are shaded red with a darker color indicating higher number of tested experiments. Protein pairs shown in white were not abundant enough to be considered ‘tested’.

If orientation is not considered (simply looking at the protein pair being covered in any orientation), 1,575 protein pairs are covered well enough to be considered tested out of a possible 1,596 protein pairs (98.7%) [total test space = (57 proteins x 56 proteins)/2; (self-pairs not considered; see below)]. Only 21 protein pairs were either never seen during sequencing or were seen in only 1 experiment (which does not meet our criteria for being ‘tested’) and are represented by a white box in both orientations (Fig. 3). Further method optimization, such as adding an additional experiment (i.e., four experimental replicates instead of three), will likely improve the number of tested combinations.

### Identification of auto activators

All experiments have internal positive and negative controls to ensure that selection conditions have worked, and sequencing was able to detect the interactions in each replicate. However, it is important to determine if any protein(s) activate the system on their own (auto-activators). This means a protein tested against a frameshifted fragment that encodes a very short random peptide showing similar growth levels passing criteria used to define an interaction in a single orientation (fused to DBD or AD). With these criteria we did not identify any protein which triggered an autoactivation of the system in a single orientation.

It is readily apparent from the interaction map (Fig. 3) that several proteins have many interaction partners. Proteins such as LMNA, LMNB1, GTF2F1, IFIT1 and SLC22A15 interact with frameshifted fragments in both orientations in multiple experiments. These proteins are annotated as ‘sticky’ in our analysis (Supp. Table 1). Using this approach, distinguishing between a “sticky” protein and an auto-activator is not always possible. Therefore, novel interactions annotated as ‘sticky’ should be viewed cautiously and warrant secondary experimental validation.

### Known and novel interactions recovered

In our strictest requirements, an interaction must be recovered in 2 or more experiments in a single orientation. A protein-protein pair interacting in one experiment in one orientation is only considered if the other orientation also interacted in at least two experiments. With these criteria, we recovered 159 full-length protein-protein interactions. Eight of these interactions are known human interacting pairs (expected from this set of PRS proteins), seven additional interactions are reported in the APID (30, 31) (but not part of the PRS), and the remaining 144 interactions are not documented in other interaction databases and are potentially novel – several with support from StringDB (32). Furthermore, 121 of 144 potentially novel interactions are recovered in both orientations and in 2 or more experiments indicating a robust, repeatable interaction. Of the 15 APID reported interactions we recovered, 13 were recovered in both orientations and 2 or more experiments.

Five of the eight PRS pairs recovered were in both orientations, and all three experiments had log_2_FC (maximum fold change in log_2_) values ranging from 1.5 to 6.2. The PPP3CA|PPP3R1 interaction was one of three PRS interactions recovered in 5 mM 3-AT, the most stringent growth conditions. Interestingly, other methods did not recover this interaction despite the interaction domains being well documented and present (12). Further investigation is needed as to why the AVA-Seq method was able to recover this interaction while no other robust method could.

We were unable to identify full length self-pairs confidently. This is likely due to hairpins forming during PCR amplification or NGS library creation. For this reason, we have removed all tested and interaction data for self-pairs to prevent any misinterpretation or error.

### Correlation between high-throughput AVA-Seq and low-throughput AVA

A key question is whether the AVA-Seq results can correlate the log_2_FC of a protein pair in read counts within a complex mixture of tested interactions to the growth of a single interaction pair, as measured by OD_600_. With the data in Fig. 1 (a low-throughput measure of liquid growth via OD_600_), the protein pairs can be detected as interacting when tested in isolation possibly due to reduced competition or crowding of neighboring cells. The experiments where NGS libraries were generated were a mixed population of 1,000’s of protein-protein pairs being tested. This means in each 5 mL culture tube, there could be 3,192 DNA fusions represented in millions of cells (100x coverage of possible combinations). In the presence of 3-AT, ‘stronger’ interactions can grow and become a higher percentage of the final DNA product – represented as more reads. Figure 4 compares the growth of individual pairs (OD_600_) to the number of read counts (maximum log_2_FC). The higher the individual growth, as measured by OD_600,_ the more likely the interaction is detected in a complex mixture (AVA-Seq).

**Figure 4.**
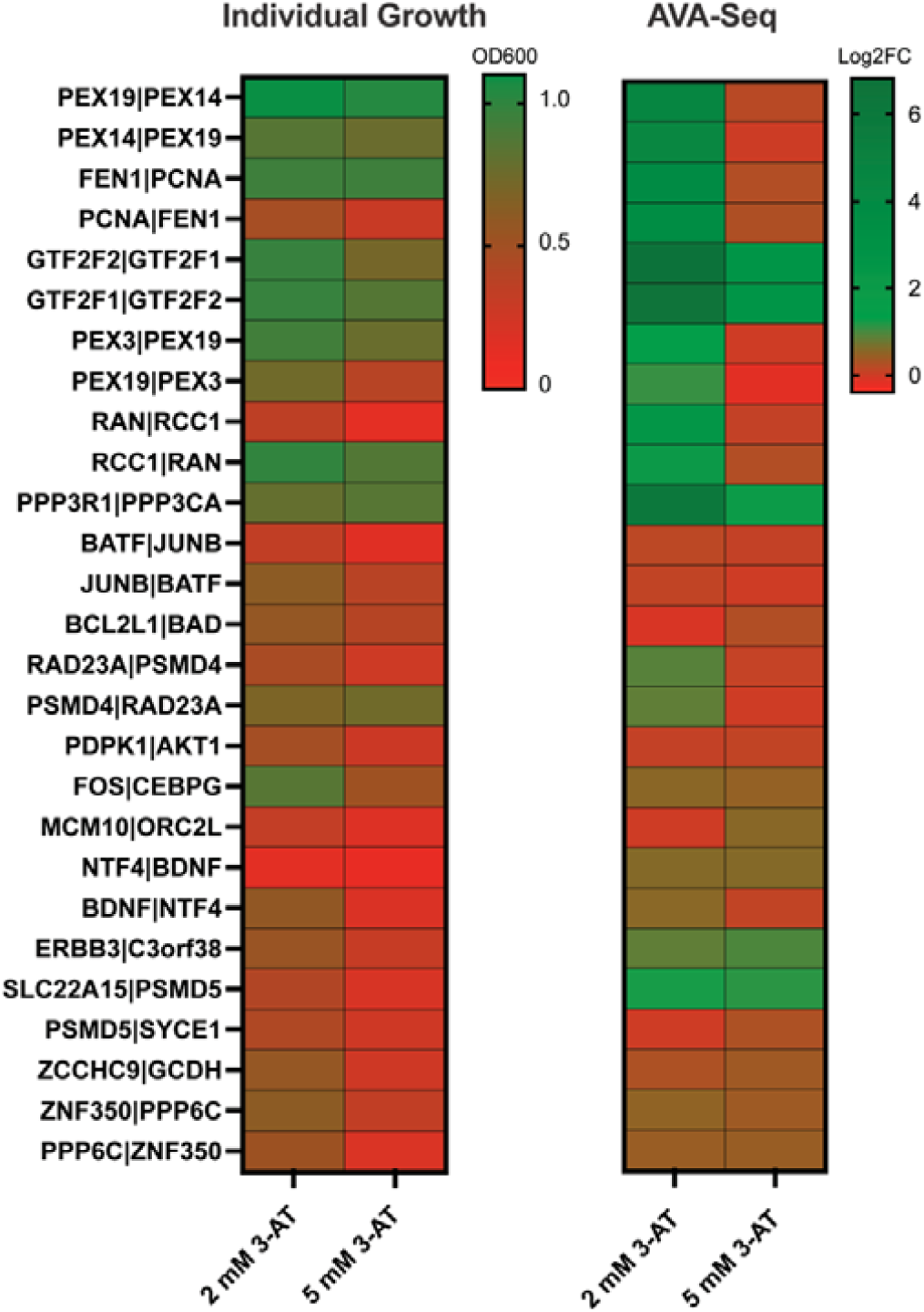
Correlation of low-throughput (individual growth in liquid culture) and high-throughput all-vs.-all sequencing (AVA-Seq). Log_2_FC max is shown for 2 mM and 5 mM 3-AT growth conditions. FDR < 0.1. Interactions were considered if the log_2_FC >0.8 and in two experiments. PRS pairs from this study are listed in both orientations in the left column. Not all orientations were successfully clones resulting in missing data for the individual growth. Selected individual RRS pairs (as reported in HsRRS-V2 (12)) ERBB3|C3orf38, SLC22A15|PSMD5, ZCCHC9|GCDH, ZNF350|PPP6C, PPP6C|ZNF350 were tested in the individual liquid growth (in those specific orientations only) and AVA-Seq high throughput. ERBB3|C3orf38 and SLC22A15|PSMD5 showed AVA-Seq growth in only 1 experiment out of three tested so they do not meet the criteria for being considered an interaction. C3orf38 and SLC22A15 are annotated as ‘sticky’ proteins which may explain their growth.

Expected interactions such as PEX14|PEX19, PEX3|PEX19, RAN|RCC1 and FEN1|PCNA show strong growth in 5 mM 3-AT conditions using the low-throughput growth experiment (Fig. 1, Fig. 4). However, although these pairs are present in the AVA-Seq (high-throughput NGS experiments), they do not show a significant increase in read counts in 5 mM 3-AT conditions indicating they do not interact in these experimental conditions. The RAN|RCC1 interaction showed higher growth in one orientation using the low-throughput method, but both DBD and AD fusions had similar log_2_FC in the high-throughput AVA-Seq experiment (Fig. 4). Competition may happen when the protein pairs are pooled together, leaving some PPIs at a disadvantage. This means we see specific pairs that grow in the low-throughput method and on solid agar (not shown) but not in the high-throughput AVA-Seq. The reason(s) remains unclear. It may be ‘simply’ one of the features of AVA-Seq (or pooled interaction methods), and it further reiterates that multiple methods can give different results with minimal overlap. Overall, our data suggests using sequencing read counts to quantify PPIs is appropriate, as shown previously in the short-read versions of the AVA-Seq system (8, 25, 26).

## DISCUSSION

We have applied the AVA-Seq protein-protein interaction method to full-length proteins using advances in synthetic biology and long-read sequencing technology. The extended dynamic range from sequencing on ONT makes applying an all-vs.-all protein interactome approach valuable – meaning the progressively weaker interactions can be detected by deeper sequencing to increase sensitivity. The AVA-Seq method takes advantage of the foundation of the B2H system and combines it with high-throughput sequencing methods. The more robust growth of a protein pair indicates a greater ability to overcome the 3-AT inhibition and correlates to a stronger protein interaction. At the same time, lack of growth in 2 mM 3-AT indicates an interaction that cannot overcome this mild inhibition. Our system challenges protein-protein interactions in 2 and 5 mM 3-AT concentrations allowing for greater sensitivity in recovering interactions that would likely be missed by other overly stringent methods (9). Furthermore, making the requirement that interactions must be captured in multiple experiments further reduces the likelihood these interactions are false.

The convergent fusion design of the pAVA plasmid in combination with NGS is central to our ability to obtain quantifiable results with AVA-Seq. Others have suggested a challenge in comparing one interaction pair to another using a traditional two-plasmid B2H platform due to different expression levels of the plasmids (i.e., low copy pBT and high copy pTRG). Yet another advantage of AVA-Seq over traditional two-hybrid systems is the DNA from both genes are contained in a single DNA fragment. This means that the copies of each gene in the pair being tested are equal. Therefore, there is no need to manipulate the gene transcription levels or adjust the IPTG concentrations using our AVA-Seq method (33).

### Perspective and Future Challenges

There are many advantages to utilizing the Oxford Nanopore platform for studying protein-protein interactions. A top benefit is the portability of the instrumentation and low cost, allowing many more laboratories to utilize the technology. Using long-read sequencing technologies will enable researchers to screen full-length protein pairs at scale without fragmentation. Individual constructs are not required; instead, an entire pool of proteins can be created and screened all-vs.-all. Additionally, utilizing full-length genes are more likely to result in biologically relevant interactions as they consider the 3D structure, multiple binding sites, etc. With the increased availability and reduced cost of synthetic DNA, we see this as a viable option for multiple sizes of projects, significantly reducing the need to first PCR and clone genes prior to protein interaction mapping. Here, utilizing full-length synthesized genes makes the entire protein pair matrix readily attainable - allowing us to increase the search space and interaction coverage (4). Compared to Illumina platforms, ONT has a higher error rate, potentially making distinguishing between a wildtype and a single amino acid variant sequence challenging. However, we are working to overcome this by utilizing unique barcodes for each protein or relying on individual codon optimization sequences (unpublished work).

It is undoubtedly apparent that no single method is the end-all and be-all of protein interaction detection. Even within our AVA-Seq system, we observe different interaction partners depending on the experimental approach. This doesn’t necessarily make one method superior to the other. The experimental design should be in line with the questions being asked. The use of the fragments allows for interactions to be localized to regions of a protein, while full-length proteins allow for more rapid screening.

### Current limitations of AVA-Seq

Current limitations for what can be screened using this high-throughput AVA-Seq method include the size of synthetic genes or gene fragments. Most commercial synthesis can synthesize genes up to 1.8 kb. For genes longer than 1.8 kb, tiling across the protein is possible, though the benefit of full-length protein folding is lost.

The protein-protein pairs in this study likely span a diverse kinetic range. This means some protein pairs will have a high affinity (low K_D_) for their partner while others are more transient (higher K_D_). Having the ability to extract quantitative interaction strengths in bulk would be extremely useful and allow us to potentially explain the difference in interactions recovered using fragment-based AVA-Seq and full-length AVA-Seq – although this is beyond the scope of this study, but is an avenue we hope to explore.

The inability to recover specific known protein-protein interactions could be due to many reasons not unique to our system: cellular localization, media composition, post-translational modification dependence, and the geometry of the fusion pair (35). Using an alternative bacterial strain could improve the recovery of specific interactions (11). Further, flow cell-based sequencing methods tend to prefer shorter pairs. This makes it more challenging when attempting an all-vs.-all interactome using a dynamic range of gene lengths. The experimental design could be further simplified upon improvement of the sequencing technologies/chemistries.

### Breakthrough applications

We demonstrate the significance of this method by recovering protein-protein interactions between a wide range of gene pairs without the need for fragmentation. We envision several trajectories for the AVA-Seq and ONT partnership. First is utilizing a pool of long protein fragments against themselves. This could be human vs. human, pathogen vs. pathogen, or host vs. pathogen interaction networks. A second application would be a more targeted interaction screening utilizing a single protein and short fragments of a known interactor or set of interactors (i.e., pathway-specific or class of enzymes). When considering the all-vs.-all experimental approach, the AVA-Seq method makes it easy to identify auto-activating proteins by simply including frameshifted fragments (which code for random peptides) in the gene pool.

The AVA-Seq method is a powerful tool to detect protein fragment-fragment interactions (8, 25, 29) and full-length protein-protein interactions. Here, we showed a novel application combining the AVA-Seq system and Oxford Nanopore Technologies MinION platform to detect full-length protein interactomes. AVA-Seq affords a dynamic range of PPIs using this long-read technology with direct readout of interactions based on the read count. These interactions are easily scalable and achievable with a single investigator.

In summary, conducting full-length PPIs with AVA-Seq is now possible, allowing researchers to move beyond colony formation. Utilizing NGS platforms allows for quantitative readout of PPIs as the log_2_FC from the high-throughput experiments correlates with the growth in individual screening using liquid culture. NGS allows for increased sensitivity in detecting weak or transient protein-protein interactions often missed with mainstream platforms. Our AVA-Seq method is an important tool in the high-throughput protein interaction mapping repertoire as it recovers a unique population of interactions. Furthermore, this manuscript demonstrates a novel application of ONT in identifying full-length protein-protein interactions.

## DATA AVAILABILITY

The data underlying this article are available xxx *ID XXXX*.

## SUPPLEMENTARY DATA

Supplementary Data are available at xxx online.

## ACKNOWLEDGEMENTS

We would like to thank the members of the Genomics Core at Weill Cornell Medicine – Qatar.

