## Supplemental Fig. 1 for "Long-read sequencing to detect full-length protein-protein interactions"

Supplemental Figure 1. A 9- and 16-hour growth comparison of selected known human interactions. Three protein pairs were tested in both fusion orientations and final OD_600_ were recorded at 9- and 16-hours in minimal media supplemented with 2 and 5 mM 3-AT . Most pairs show higher growth in both 2 and 5 mM 3-AT at the 9 hour time point.

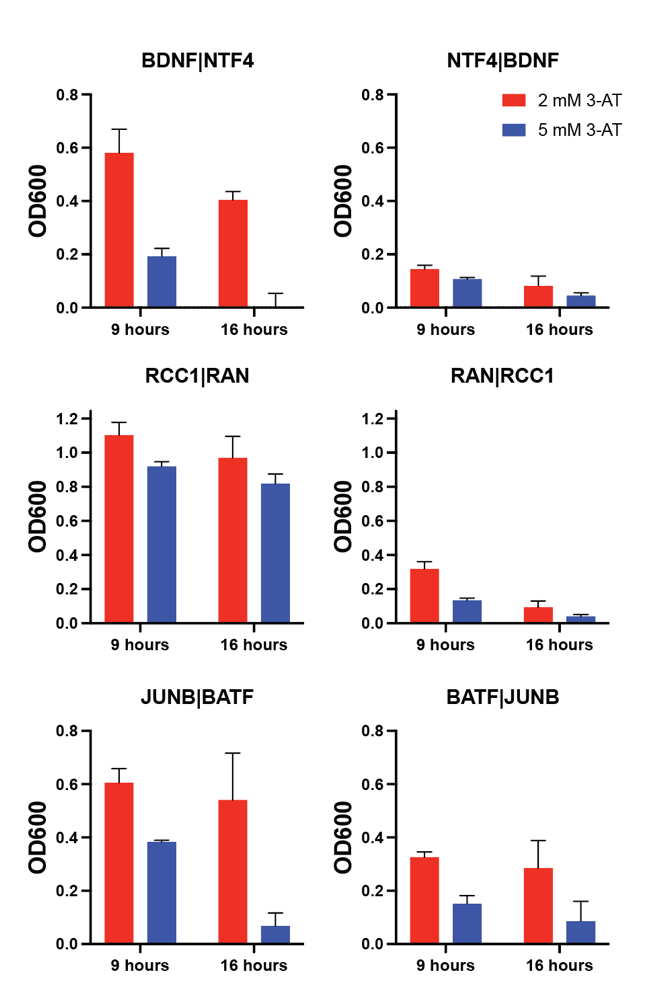
